# Metadata Harmonization from Biological Datasets with Language Models

**DOI:** 10.1101/2025.01.15.633281

**Authors:** Alexander Verbitsky, Patrick Boutet, Mohammed Eslami

## Abstract

Biomedical research faces significant challenges in harmonizing metadata across diverse datasets due to inconsistent labeling and the lack of universally adopted ontologies. Conventional solutions, such as Common Data Elements, face adoption difficulties as they impede scientific progress by requiring researchers to navigate through thousands of standardized terms with subtle variations. Tools such as laboratory information management systems, while designed to enforce standardization, can hinder research progress when their rigid standards conflict with domain-specific documentation needs and evolving research practices. As a result of these challenges, researchers maintain their own annotation systems, leading to disconnected datasets that are difficult to integrate across studies.

This study presents a novel approach using large language models to automatically standardize researcher annotations to standards within ontologies. The approach is applied to multiple domains such as oncology, alcohol research, and infectious disease. Data augmentation strategies are presented to align training data with the space of human representations. These strategies generate realistic variations of standard terms to simulate how researchers naturally document their work, especially valuable in domains lacking the extensive terminology mappings needed for training language models. Experiments with fine-tuned GPT-2 variants show up to 96% accuracy on in-dictionary tasks and 17% on out-of-dictionary tasks, outperforming traditional techniques and zero-shot GPT-4o applications. This implies that there can be up to a 96% reduction in metadata standardization labor if a term exists in an ontology. We also show a significant trade-off between domain-specific models versus those that aim to generalize across domains such as infectious disease or alcohol research. While larger models excel at generalization, fine-tuned models consistently outperform on domain-specific terminology. This approach enables more efficient and accurate research data integration across biomedical fields, though out-of-dictionary generalization remains a challenge across all model sizes.

## Introduction

Artificial intelligence (AI) models require a large, curated training corpus. While data generated across a variety of biomedical research domains is only increasing, they are not curated in a way where they can be rapidly used to train AI models ^1^. Inconsistencies in data annotations primarily arise because the data comes from multiple sources and modalities (e.g. genomics, transcriptomics, proteomics), institutions, and researchers. Ontologies, controlled vocabularies, and common data elements are among the tools that seek to ensure data annotations are consistent across all sources. These tools can ensure data are annotated and connected in a consistent, standardized manner. There is no shortage of them in biomedical research from synthetic biology ^2,3^, to disease specific ontologies such as those in cancer research ^4,5^ and infectious disease ^6,7^, to even foundations setup for protocol ^8^ and clinical trials ^9,10^ . Yet, data uploaded across these domains are disconnected from these tools. Therefore, the same annotations are represented differently by researchers and labs ^1,11^. Due to this diversity, data harmonization for downstream integration with other datasets or tools typically takes over 40% of an analyst’s time ^12–14^.

State-of-the-art technologies to bring the ontologies to the researcher are primarily electronic lab notebooks ^15^, or customized webforms ^16,17^. These technologies significantly hinder research progress by overwhelming users with drop-down selections before even running an experiment. Natural language processing and machine learning have been used to support standardization ^18–20^. However, these techniques require extensive surrounding text, such as full paragraphs from publications or abstracts, and cannot be directly applied to annotations in filenames or spreadsheets where there is minimal context.

This calls for new, innovative approaches to standardize annotations across data sources while minimizing impediments to a user’s workflow. These new approaches should be able to automatically standardize terms to a specified ontology once a variant is observed with minimal context. In other words, we want to minimize the amount of information a researcher would need to provide every time a new dataset needs to be standardized. We present a study that uses language models to support this standardization. We first show the potential of the use of language models to standardize annotations in cancer research, one of the most well-funded domains in biomedical research. There are ontologies, common data elements, and even a collected set of representations that researchers have used when annotating their own datasets. In this effort, we show the language models can reduce the labor of standardization >96% if the term being represented is within the ontology. But, it can only achieve 17% reduction if the term is not within the ontology. We then move to a domain whose metadata are not as well curated, alcohol research, and present a technique to train language models for standardization. We show that domain-specific, small language models can nearly automate the task for small, lab-specific controlled vocabularies. These small models, however, will never be able to resolve a term outside of those controlled vocabularies. Finally, with the generalization capabilities of large language models, we wanted to see if the standardization models were transferable across domains. We find that fine-tuned, domain-specific models significantly outperform generalized models.

## Results

### Cancer Ontology Data Collection

Cancer is one of the most studied domains in biomedical research and as a result has no shortage of data or ontologies. We took advantage of a training corpus that maps malformed researcher annotations from data, or synonyms, to the standard term (or command data element) within an ontology. We combined terms from multiple sources, including the National Cancer Institute Thesaurus (NCIt) ^21^, Cancer Data Standards Registry and Repository (caDSR) ^22^, Genomic Data Commons (GDC) ^23^, International Classification of Diseases for Oncology (ICD-O3) ^24^, and Medical Dictionary for Regulatory Activities (MedDRA) ^25^. Our compiled training dataset contains a total of 691,220 terms (Figure 1A). Most of these terms (617,978) are associated with semantic types, a NCIt classification system that categorizes terms based on their nature or role within the biomedical domain. We focused on the five largest semantic types: “*Finding*,” “*Neoplastic Process*,” “*Disease or Syndrome*,” “*Laboratory Procedure*,” and “*Quantitative Concept*.” This subset, comprising 165,255 terms, not only represents the most populous groups within the dataset, but also provides a more manageable scope for the harmonization task. Chemical names and procedural semantic types were excluded (specifically, “*Pharmacologic Substance*”, “*Organic Chemical*”, “*Therapeutic or Preventive Procedure*”, and “*Intellectual Product*”) to concentrate on terms that follow more typical natural language structures rather than technical jargon. Additionally, 154,432 ambiguous term mappings, instances where a single representation maps to multiple target standards, were identified. Only a single one of these mappings was included in the training corpus to prevent ill-formed data points. The rest were held out to be appended after the model’s prediction in a post-processing step to facilitate named entity disambiguation.

**Figure 1.**
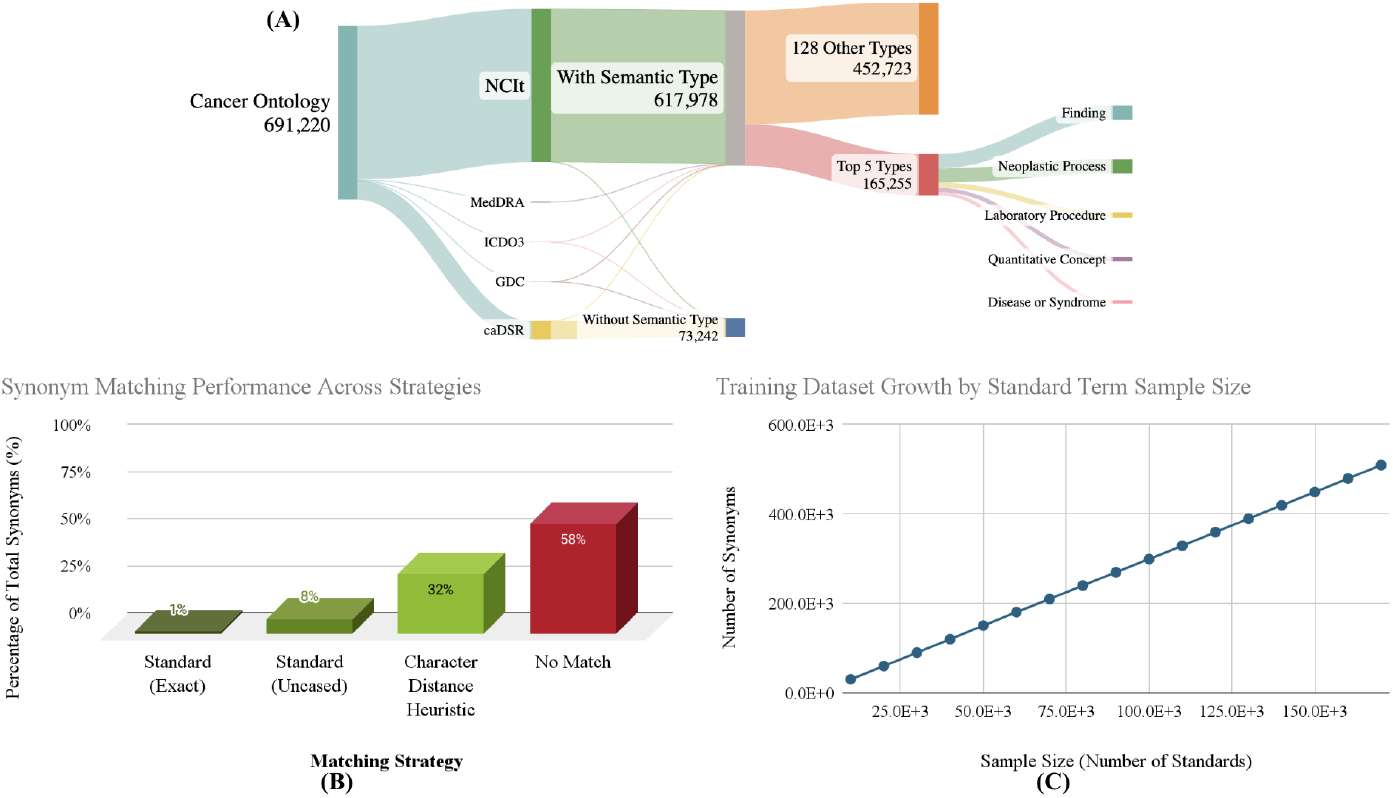
**(A)** Composition of the cancer ontology dataset. Distribution of terms across categories, including those with semantic types (617,978), the subset of top five semantic types used for fine-tuning (165,255), and terms without semantic types (73,242). **(B)** Synonym matching performance across different strategies for 434,841 synonyms matched against 259,707 unique standards from our compiled cancer ontology. Standard (Exact): exact string matching; Standard (Uncased): case-insensitive matching; Character Distance Heuristic: fuzzy matching based on character differences; No Match: synonyms unmatched by any method. 58% of synonyms remained unmatched, demonstrating limitations of basic matching. **(C)** Fine-tuning dataset growth as a function of standard term sample size, showing a near-linear relationship between the number of standard terms included and the total number of synonyms in the dataset.

With this dataset, we wanted to assess the scale of the harmonization problem. Namely, can simple methods, such as capitalizations, direct matches to previous synonyms, or string distances solve the standardization problem (Figure 1B). In this context, a synonym is a malformed representation of a standard term that conveys the same or similar meaning. Synonyms can take various forms, including acronyms (e.g., “MRI” for “Magnetic Resonance Imaging”), abbreviations (e.g., “Ca.” for “Cancer”), lexical variants (e.g., “tumor” and “tumour”), word reorderings (e.g., “Cancer of the Lung” for “Lung Cancer”), or conceptually related terms (e.g., “Neoplasm” for “Tumor”). To simulate a real-world scenario where the association between synonyms and their standards is unknown, we tested all synonyms (434,841) against all standards (259,707) in our cancer ontology using various matching techniques. Standard exact matching, requiring perfect string alignment, successfully matches only a small fraction of terms. Even case-insensitive matching (Standard Uncased) yields only marginal improvement. A more sophisticated approach using an indel distance heuristic performs better, but still leaves a significant portion of terms unmatched. 58% of synonyms in our dataset remain unmatched using these conventional techniques. Moreover, heuristic approaches are limited to known standards, necessitating more advanced harmonization methods such as generative models which can potentially standardize unfamiliar terms.

To develop our AI-driven approach, we constructed a fine-tuning dataset consisting of 141,065 terms, including 83,904 synonyms of 57,161 standards. This dataset provides a diverse range of term variations and relationships for training our models. We observed a near-linear relationship between the number of standard terms and the total number of synonyms in our training dataset (Figure 1C), indicating that, on average, each standard is associated with three synonyms. This results in a uniform density of term variations across the vocabulary. For evaluation, separate validation and test sets were created, each containing 200 terms. The validation set guided early stopping, tracked performance, and identified the best model during training. The test set was used to assess the final model’s performance.

### Large Language Models for Cancer Metadata Harmonization

Language models have the potential to map malformed representations of standards to their standard terms within an ontology ^26–28^. Our focus here with the ubiquity of large language models (LLMs) is to see if this mapping can occur with minimal context. LLMs are trained on vast amounts of data and so already should account for much of the context one would provide. They can be used in two ways: 1) fine-tuned with a set of malformed representations, or synonyms, that should be mapped to standards ^28^. 2) zero shot, where we simply provide a prompt and ask for a structured, parseable output ^26^. We tested both uses of LLMs.

LLMs were fine-tuned context-free on a curated cancer ontology dataset using the prompt *The standardized form of “synonym” is “standard”*. During fine-tuning, the model learns to predict the correct “standard” given the “synonym” and preceding text. This allows the model to learn the associations between synonyms and their standardized forms. During testing and application, the prompt was modified to *The standardized form of “synonym” is “*, creating a completion task. This approach challenges the model to generate the standardized term based on learned associations, applying the knowledge it acquired during fine-tuning. The resulting system automates harmonization while allowing researchers to continue using their preferred terminology, alleviating the burden of manual standardization.

GPT-2 was selected for this study due to its computational efficiency, well-documented fine-tuning process, and unrestricted commercial use. While newer, larger models could potentially offer better out-of-dictionary (OOD) performance, GPT-2 provides an efficient balance of performance and practicality. Five iterations of four different-sized GPT-2 model variants (GPT-2, GPT-2 Medium, GPT-2 Large, and GPT-2 XL) were fine-tuned and evaluated using in-dictionary (ID) and OOD accuracy metrics. All models were trained with an early stopping criteria, halting training after 20 consecutive evaluations without validation accuracy improvement at 0.05 epoch increments. Accuracy was measured with no credit given to partial matches, providing the most stringent assessment of model performance. ID accuracy evaluates the model’s ability to standardize new synonyms of standards present in the fine-tuning dataset, while OOD accuracy assesses its capacity to generalize to entirely new standards and their synonyms that the model has never seen before. The OOD test was made even more challenging by exclusively including terms from semantic types not encountered during training.

As expected, larger models generally performed better, with GPT-2 Large achieving the highest accuracies: 96% for ID and 17% for OOD tests. Interestingly, the largest model, GPT-2 XL, performed slightly worse than GPT-2 Large (95% ID, 15% OOD). The substantial gap between ID and OOD performance (78-79% across models) underscores the challenge of generalizing to new concepts and semantic types in specialized domains. While models effectively standardize variations of familiar terms, they struggle with novel standards and synonyms, particularly from unseen semantic categories, highlighting the complexity of true out-of-domain generalization in biomedical nomenclature.

Given the requirement of fine-tuning, we also analyzed the relationship between model performance and the cost of fine-tuning (Figure 2C). GPT-2 Large achieved the highest ID accuracy at 96% with a training cost of $29, outperforming GPT-2 XL, which reached 95% accuracy at $143. Despite having fewer parameters (774M vs 1.5B), GPT-2 Large delivered better performance at a fraction of the cost, making it the more cost-effective choice. Ultimately, the task here is so specific that it does not require all of the capabilities of larger LLMs.

**Figure 2:**
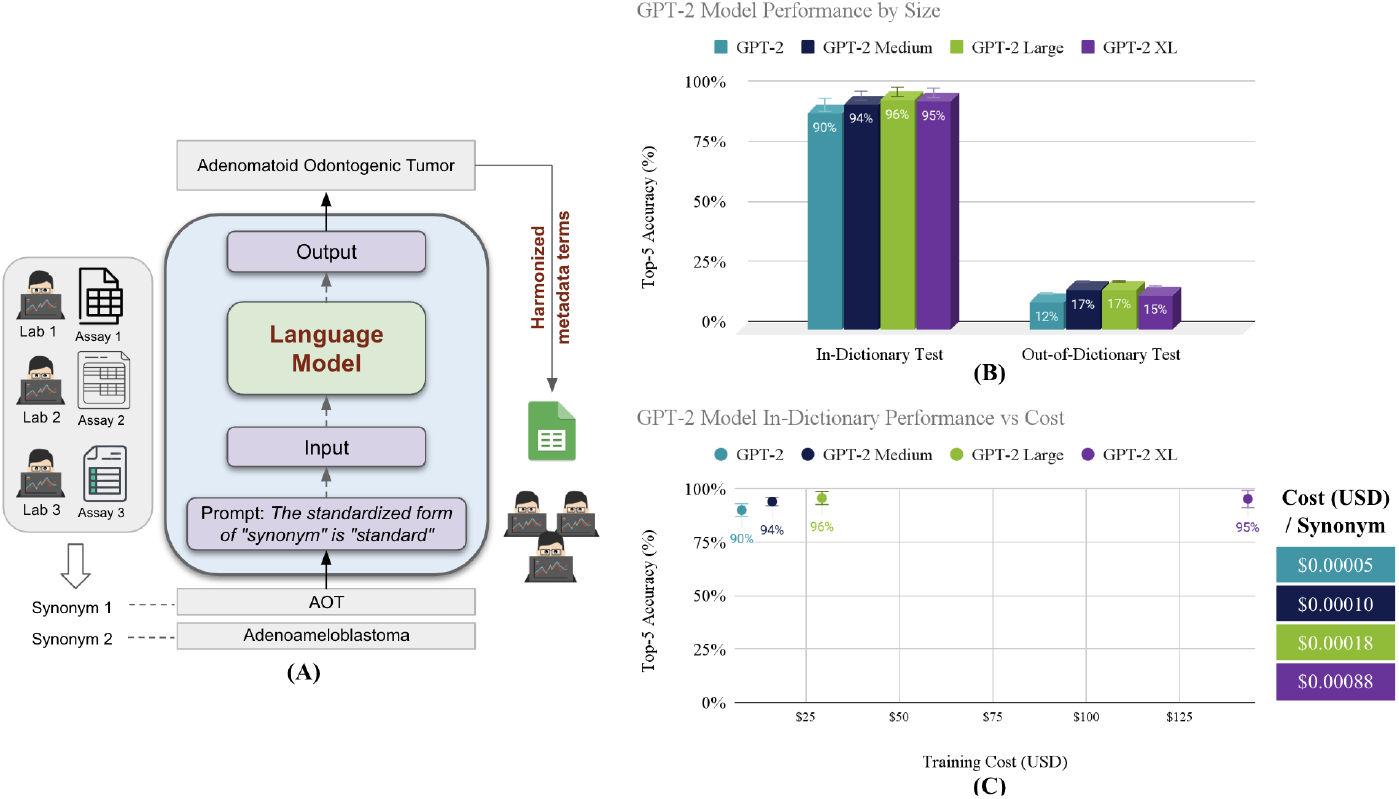
**(A)** The LLM-based harmonization system. The model takes a synonym as input, along with a prompt specifying the task, and outputs the standardized form of the term. **(B)** GPT-2 model top-5 performance by size for in-dictionary and out-of-dictionary cancer metadata harmonization tests. In-dictionary accuracy ranges from 90% to 96%, while out-of-dictionary accuracy is 12% to 17%. **(C)** GPT-2 model in-dictionary performance vs. training cost. GPT-2 Large achieved 96% accuracy at $29, while GPT-2 XL reached 95% accuracy at $143.

For the zero-shot use case, we benchmarked the same system with GPT-4o (gpt-4o-2024-08-06), OpenAI’s most recently released model at the time of publication. Despite GPT-4o’s broader generalization capabilities due to its larger parameter size and extensive pre-training, it underperformed compared to our fine-tuned GPT-2 models on familiar terminology. GPT-4o achieved an average of 25% top-5 accuracy (±5%) across all tests, modestly improving on our models’ 17% OOD accuracy, but falling far short of our models’ ≥90% accuracy on familiar ID terms. GPT-4o’s slight improvement in handling synonyms of standards unfamiliar to our models reflects its broader exposure to varied linguistic contexts, but it lacks the fine-tuned specialization required for high accuracy on known standards. Our results suggest that while large, general-purpose models such as GPT-4o can contribute to improved performance on novel terms, they are not sufficient alone to solve the harmonization task. Domain-specific, adaptive models remain essential for tasks such as cancer-related metadata standardization, where familiarity with the field’s specific nuances significantly boosts performance.

### Harmonization of Terms Where Training Sets Do Not Exist

As previously mentioned, cancer research has had the opportunity to develop standards, but also curate common malformations, or synonyms, that have been actively used to represent standards. This is not true in other domains such as addiction research. Given our experience with alcohol research, we applied our technology to this domain, where a curated set of synonyms mapping to standards does not yet exist.

We compiled a comprehensive alcohol ontology of 27,499 standards from authoritative sources, including the World Health Organization (WHO) Lexicon of Alcohol and Drug Terms (141) ^29^, the Health Research Board (HRB) National Drugs Library (550) ^30^, and the consensus measures for Phenotypes and eXposures (PhenX) Toolkit (26,808) ^31^(Figure 3A). Unlike the cancer ontology, which includes multiple synonyms per standard, this ontology contains only standardized terminology without synonyms. This presents a challenge for training harmonization models, not only in the alcohol domain, but in any specialized field with similar limitations.

**Figure 3:**
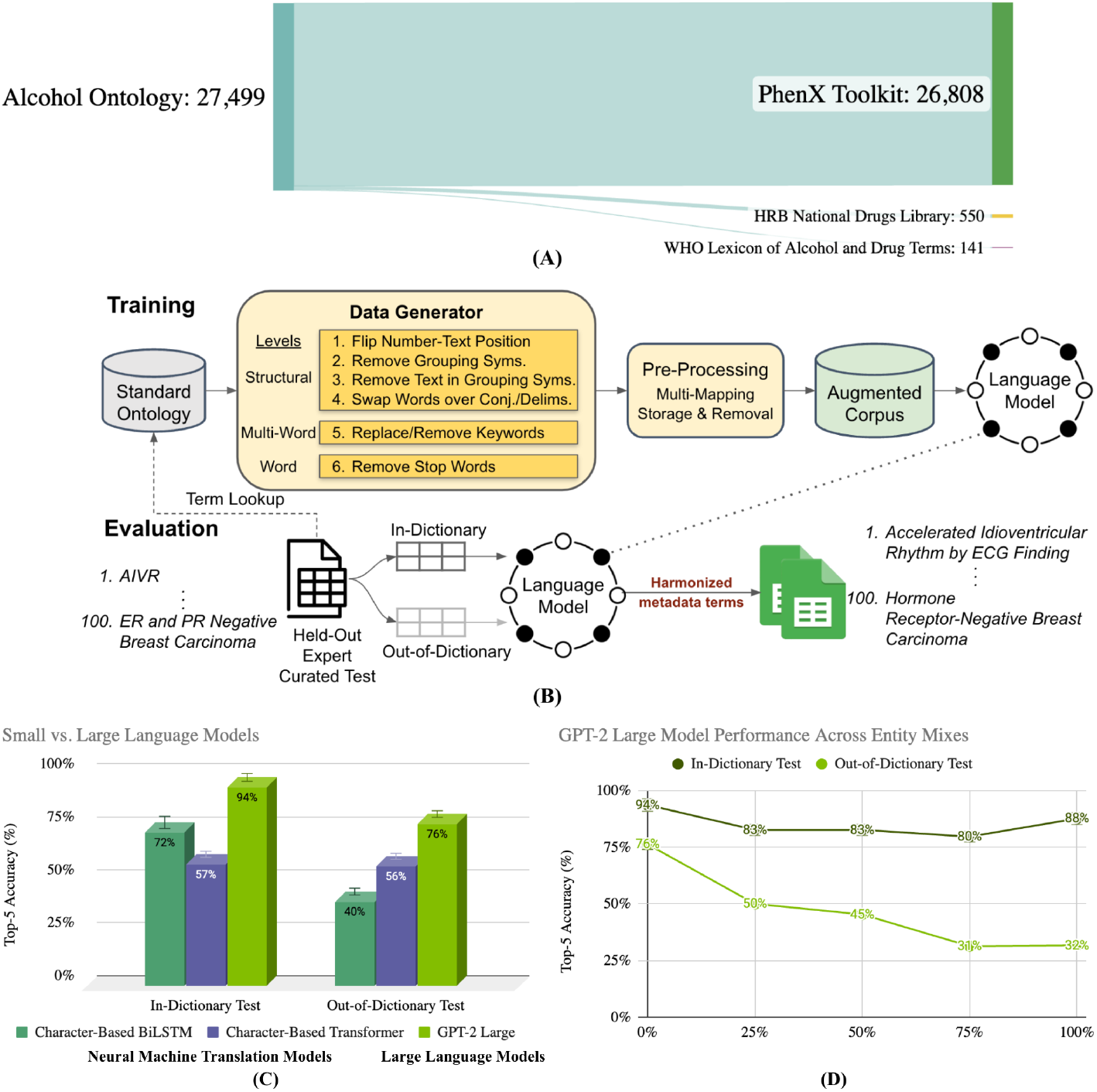
**(A)** The distribution of terms across standard sources in the combined alcohol ontology. The alcohol dataset includes 27,499 standard terms, mainly from the PhenX Toolkit. **(B)** Language model training and evaluation process for metadata harmonization. Standard ontology terms flow through data generation and pre-processing to create an augmented corpus, which is used to train language models. Models are then evaluated on in-dictionary and out-of-dictionary tests. **(C)** Top-5 accuracy comparison of small vs. large language models on alcohol terminology harmonization, showing in-dictionary and out-of-dictionary test results for NMT models (Character-Based BiLSTM and Transformer) and LLMs (GPT-2 Large). **(D)** GPT-2 Large model performance on alcohol terms, with training conducted on varying mixes of alcohol and bacteria terminologies. ID and OOD test accuracies decline as the proportion of bacteria terms (Entity 2) in the training mix increases from 0% to 100%.

With only the standards available, we synthetically generated a set of data that would create synonyms of those standards (Figure 3B). This synthetic training dataset would serve as the training data for a harmonization model. Six transformations of three distinct types were applied to the standards (details can be found in the Methods section). The pre-processing module then handles multi-mapping instances and removes conflicting variations as we discussed with the cancer ontology. This approach created a training corpus of over 200K synonym-to-standard mappings that can be used to train a model.

We trained two types of language models - Small Neural Machine Translation (NMT) models and LLMs - on the augmented terminology corpus and evaluated their harmonization performance (Figure 3C). Results highlight a trade-off between domain-specific accuracy and generalization. Among the small NMT models, the Character-Based Bidirectional Long-Short Term Memory (BiLSTM) achieved the highest ID accuracy at 98%, followed by the Character-Based Transformer at 50%. In contrast, LLMs such as GPT-2 and GPT-2 Medium achieved 87% and 92% ID accuracy, respectively. However, in OOD tests, LLMs demonstrated significantly better generalization. GPT-2 Medium reached 51% OOD accuracy, surpassing both the Character-Based BiLSTM’s 3% and the Character-Based Transformer’s 24%. This performance disparity illustrates the complementary strengths of small, specialized models and larger models that excel in broader generalization tasks.

Finally, to further explore model generalizability and domain adaptation capabilities, as we did with different semantic types in cancer, we expanded our investigation to microbiology, specifically bacteria names, building on our past performance ^32,33^ in the domain as well as the rising concern of antimicrobial resistant (AMR) bacteria. Given the growth in funding to study AMR bacteria, with over $8 billion invested globally between 2008 and 2030 ^34^, we anticipate the generation of hundreds of thousands of datasets, each contributing additional taxonomic representations to the research landscape. Therefore, we curated 627,937 standard names of bacteria drawn primarily from the National Center for Biotechnology Information (NCBI) Taxonomy (627,793) ^3^ with contributions from our prior work ^32^ with National Institute of Standards and Technology (NIST) (144). Both ontologies share the characteristic of containing only standard terms, setting the stage for investigating harmonization techniques in fields where synonym diversity is limited and exploring the potential for cross-domain harmonization. Next, an entity mixing experiment was conducted by blending alcohol and bacteria terminology (Figure 3D). As the proportion of bacteria terms (Entity 2) increased from 0% to 100%, a steady decline in the GPT-2 Medium model’s performance was observed in both ID and OOD tests. ID accuracy decreased from 92% to 65%, while OOD accuracy dropped from 51% to 13%. This decline highlights the difficulty of maintaining accuracy across diverse domains even with synthetically generated data. This suggests that domain-specific models may be required for achieving optimal harmonization in specialized fields.

## Discussion

An equivalent of a “drag and drop” curation process for biomedical research will significantly streamline the application of AI in the domain. We showed that language models can be used in a variety of ways (fine-tuning vs zero-shot) to achieve semi-automated curation. Traditional heuristic methods, such as fuzzy matching, perform well for in-dictionary terms but struggle with out-of-dictionary variations, limiting their effectiveness in real-world scenarios. Small NMT models, such as character-based BiLSTMs, offer improvements over heuristics, demonstrating non-zero OOD performance. These models excel in specialized tasks within a narrow context, but lack the broad knowledge necessary for adapting to novel terms or concepts. Medium to large-sized models, exemplified by GPT-2, strike a balance between specificity and generalization, offering improved OOD performance while maintaining high ID accuracy. At the far end of the spectrum, very large language models, such as GPT-4o, provide superior generalization capabilities, addressing a wider range of harmonization tasks across multiple domains. However, these models come with increased computational costs and potential overgeneralization, which may introduce errors in highly specialized fields. For instance, large language models benefit from a general understanding of English, which is particularly advantageous in domains such as cancer and alcohol research, where many terms overlap with everyday language. This pre-existing knowledge base contributes to their strong performance in harmonizing representations of non-domain specific words. However, in fields such as microbiology, where bacterial names are less common in general discourse, the advantage of larger models is less pronounced.

Throughout our work, particularly with the augmentation strategies, we noticed that a training corpus of word transformations that align with the evaluation data is key to an accurate model. Beyond having a model that was rich in standards, the variations that aligned were central to achieving high standardization accuracy. This means that while data augmentation can significantly expand a training corpus through means such as extensive typo generation, this would produce an unrealistic model when deployed on real variations. This implies that even with limited initial data, creative approaches to corpus expansion can significantly enhance model performance.

The approach we presented includes no additional context beyond the variation and standard to the model. This design choice reflects the practical constraints of real-world research environments, where time and resources for data annotation are limited. However, the performance gap between ID and OOD accuracy across all model sizes suggests that additional context could improve harmonization outcomes, particularly for ambiguous terms or cross-domain applications. Future research should explore adaptive systems that enhance accuracy while maintaining contextual efficiency. These systems could employ confidence thresholds to identify cases where the model’s certainty is low, prompting a request for more contextual information from the user. This iterative approach should leverage additional context only when needed to dynamically adapt to each harmonization task’s complexity, balancing accuracy and efficiency.

Finally, the choice of model size and architecture depends on the specific requirements of the harmonization task, including the breadth of the domain, the availability of training data, and the desired balance between accuracy and generalization. While very large language models (>1 billion parameters) offer a wide range of capabilities, including image recognition and code generation, the specific task of terminology harmonization may not require this full spectrum of abilities. It is more cost-effective and practical to use a tool suited for the specific task at hand. For domains with stable, well-defined vocabularies, smaller specialized LLMs offer the best balance of accuracy and efficiency. In contrast, fields with rapidly evolving terminology or frequent encounters with novel terms benefit from larger models’ generalization capabilities, despite their slower inference speeds and higher resource demands.

## Methods

### Data Sources and Preparation

Ontologies for cancer, alcohol, and bacteria terminology were constructed from authoritative sources, each undergoing domain-specific processing. Data was exported from NCIt, caDSR, GDC, ICD-O3, and MedDRA for the cancer ontology. When merging these datasets, semantic type information from NCIt was extended across all standard items by matching texts to terms with existing semantic types. The alcohol ontology combined exported terms from PhenX Toolkit, WHO Lexicon of Alcohol and Drug Terms, and HRB National Drugs Library. A held-out annotated dataset from WCAAR was reserved for alcohol terminology evaluation. For bacteria, data exported from the NCBI Taxonomy served as the primary source, supplemented by standard conventions from NIST. The NCBI Taxonomy was processed by merging taxonomic hierarchies, scientific names, and lineage information, filtering to include only bacteria-related terms while preserving taxonomic relationships and classifications.

Common processing steps were applied across all ontologies to ensure data consistency and quality. Each source underwent standardization to create consistent standard-synonym pairs, with each standard term also appearing as its own synonym. Throughout the merging process, all original information and columns from each source were retained, including file location and source terminology, enabling traceability back to the original data.

Multi-mappings were identified, stored separately, and removed from each primary dataset to ensure data integrity for model training. These multi-mappings, where a single synonym maps to multiple standards, were identified using case-insensitive comparison and stored in a structured format, mapping each standard to its associated standards and synonyms, excluding self-references. The removal process prioritized keeping standards over synonyms. For remaining ambiguities, additional criteria were applied sequentially to create more learnable training data: preference for highest string similarity between the standard and synonym (to maintain syntactic relationships), and text with title case capitalization. This aimed to preserve the most representative examples for model training. The resulting clean, uniform structure facilitated cross-domain analysis and model development, producing comprehensive ontologies suitable for harmonization tasks.

All of the curation data can be found as supplementary data attached to this paper.

### Harmonization Heuristics

Exploratory analysis applied exact, case-insensitive, and similarity-based heuristic matching to suggest potential standards. Exact matches were used when the input text perfectly aligned with a known standard or synonym. Case-insensitive matching provided flexibility when the input text differed by capitalization. Additionally, a normalized indel similarity assessed how closely input terms matched existing standards. The indel distance calculates the minimum number of insertions and deletions required to change one sequence into another. This measure, calculated as 1 - (distance / (len1 + len2)), produces a score between 0 and 1. A threshold of 0.8 was applied to identify matches, accounting for minor variations that exact matching does not handle.

### Data Augmentation

A multi-level data augmentation approach was implemented to generate realistic term variations for ontologies lacking diverse synonyms. This approach operates at four levels: character, sub-word, word, and structural, applying functions to create synthetic synonyms that mimic real-world variations.

Character-level modifications simulate typographical errors and formatting changes through operations such as duplication, deletion, insertion, and transposition of characters, including keyboard proximity-based substitutions. Sub-word manipulations modify parts of words by removing common endings, replacing phonemes, and altering punctuation. Word-level techniques generate concise representations by removing non-essential words, replacing terms with synonyms, and creating abbreviations. Structural-level augmentations modify the organization of multi-word terms by altering numerical and textual element positions, manipulating grouping symbols, and rearranging words around conjunctions or delimiters.

The augmentation process involves selecting a subset of operations most relevant to the target domain from the full suite of functions. For example, in the case of the alcohol ontology, eight specific operations were chosen, including manipulations of number-text positions, grouping symbols (both removal and content manipulation), conjunctions, delimiters, keywords (replacement and removal), and stop words. These selected operations are then applied to existing terms, ensuring generated synonyms realistically represent expected variations while exploding the ontology. This approach creates a richer, more diverse model training dataset, addressing initial data scarcity.

### Model Architectures

LLMs and small NMT models were used for the harmonization task. Four GPT-2 variants—base (124M parameters), Medium (355M), Large (774M), and XL (1.5B)—were fine-tuned using the Hugging Face Transformers library. Two character-level NMT models, a BiLSTM network (7M) and a Transformer (550K), were implemented and trained using Keras.

The NMT models were designed to process input sequences at the character level. The BiLSTM model has an encoder-decoder architecture. The encoder comprises an embedding layer followed by a bidirectional LSTM layer, while the decoder includes an embedding layer, an LSTM layer, and a dense output layer. The encoder’s final hidden states initialize the decoder’s state, allowing the model to capture contextual information from both past and future tokens in the input sequence. The character-based Transformer adapts the standard Transformer architecture for character-level processing. It incorporates positional embeddings to encode sequence order, multi-head attention mechanisms to capture character relationships, and feed-forward neural networks in both encoder and decoder. Layer normalization and residual connections throughout the structure stabilize training and facilitate gradient flow.

### Model Training

LLMs and NMT models followed different training approaches due to their architectural differences. For LLMs, input data was formatted with the prompt *The standardized form of “synonym” is “standard”*, while NMT models were trained to directly output the harmonized form given an input term without a specific prompt structure. The cancer dataset included 141K entries, and the alcohol dataset contained 200K. The validation datasets had 200 items, while the ID and OOD test sets had 100 each.

Hyperparameter optimization used a combination of manual tuning and grid search. For LLMs, key hyperparameters included a maximum of 100 training epochs, per-device train and evaluation batch sizes of 8, and the AdamW optimizer with a learning rate scheduler employing a linear warmup followed by a linear decay. Weight decay was set to 0 to prevent over-regularization. For NMT models, hyperparameters included a maximum of 300 epochs, a batch size of 256, a latent dimensionality of 512 for the encoding space, and an embedding dimension of 64. The Transformer-based NMT model used 8 attention heads and 1 encoder/decoder layer. The BiLSTM model used the RMSprop optimizer with a learning rate of 0.001, while the Transformer model used the Adam optimizer with a Noam learning rate schedule. Both NMT models used sparse categorical cross-entropy as the loss function.

Model robustness was enhanced through dropout and a callback system implementing early stopping techniques. LLMs were evaluated every 10% of an epoch using a beam search with 5 beams, 5 return sequences, and a diversity penalty of 0.8 to encourage varied outputs. These models used a patience of 20 evaluations, monitoring validation ID accuracy for early stopping, and included callbacks for saving tokenizers alongside model checkpoints. NMT models, evaluated at the end of each epoch, employed an early stopping callback monitoring validation loss with a patience of 10 epochs and a minimum delta of 0.0001. Additional callbacks for NMT models included checkpointing and detection of NaN losses to identify potential training instability. For all architectures, the best-performing model based on their respective monitored metrics was automatically loaded at the conclusion of training.

### Evaluation of Harmonization Tasks

For the LLM models, outputs were generated using beam search with a width of five, producing the top candidate harmonizations. These candidates were then refined by cross-referencing with the multi-mappings identified during data preparation. If a match was found, all related mappings were included in the final top outputs, enriching the model’s predictions with potential alternative standardizations. Beam search was not used for the NMT models.

Model performance was evaluated using a combination of accuracy and additional metrics. ID accuracy evaluated the model’s ability to harmonize new synonyms for known standards, while OOD accuracy assessed generalization to unseen standards and synonyms. Beyond accuracy, additional metrics included BiLingual Evaluation Understudy (BLEU) score and Character Error Rate (CER). BLEU score quantifies n-gram precision between model outputs and reference standards, offering a similarity measure from 0 (no overlap) to 1 (perfect match). CER, calculated as the Levenshtein distance divided by the standard term length, measured string-level differences between model predictions and correct terms.

For benchmarking, OpenAI’s GPT-4o model (gpt-4o-2024-08-06) was evaluated zero-shot across six datasets, with five iterations per test and batches containing up to 100 terms. The following system prompt was used to guide the model:

> *You are tasked with harmonizing cancer-related terms to their corresponding standards, using your knowledge of cancer terminology. These terms span semantic types “Finding*,*” “Neoplastic Process*,*” “Disease or Syndrome*,*” “Laboratory Procedure*,*” and “Quantitative Concept*.*” Most standards come from the NCI Thesaurus*.
>
> *Prefer standards that are written out in full (i*.*e*., *no abbreviations or acronyms), unless the context absolutely requires an abbreviation*.
>
> *For each term provided, return:*
>
> 1. *The corresponding standard (i*.*e*., *the term that most closely matches the meaning and context of the input)*.
> 2. *The top 5 possible standards, ranked from most to least likely*.

The user prompt then provided the model with a list of terms to harmonize:

> *Please harmonize these terms:*
>
> *[list of terms]*

## Funding Acknowledgement

Funding for this work was provided by the Defense Advanced Research Projects Agency and the Department of Interior under Contract No. 140D6319C0029, Federal funds from the National Cancer Institute, National Institutes of Health, under Contract No. 75N91019D00024, and 75N94020C00003, as well as support from Agreement 23X098 with “the Frederick National Laboratory for Cancer Research, currently operated by Leidos Biomedical Research, Inc.. The content of this publication does not necessarily reflect the views or policies of the government, and no official endorsement should be inferred.

## Notes

### Competing Interest Statement

The authors have declared no competing interest.

